# Dynamics Robustness of Cascading Systems

**DOI:** 10.1101/071589

**Authors:** Jonathan T. Young, Tetsuhiro S. Hatakeyama, Kunihiko Kaneko

## Abstract

A most important property of biochemical systems is robustness. Static robustness, e.g., homeostasis, is the insensitivity of a state against perturbations, whereas dynamics robustness, e.g., homeorhesis, is the insensitivity of a dynamic process. In contrast to the extensively studied static robustness, dynamics robustness, i.e., how a system creates an invariant temporal profile against perturbations, is little explored despite transient dynamics being crucial for cellular fates and are reported to be robust experimentally. For example, the duration of a stimulus elicits different phenotypic responses, and signaling networks process and encode temporal information. Hence, robustness in time courses will be necessary for functional biochemical networks. Based on dynamical systems theory, we uncovered a general mechanism to achieve dynamics robustness. Using a three-stage linear signaling cascade as an example, we found that the temporal profiles and response duration post-stimulus is robust to perturbations against certain parameters. Then analyzing the linearized model, we elucidated the criteria of how such dynamics robustness emerges in signaling networks. We found that changes in the upstream modules are masked in the cascade, and that the response duration is mainly controlled by the rate-limiting module and organization of the cascade's kinetics. Specifically, we found two necessary conditions for dynamics robustness in signaling cascades: 1) Constraint on the rate-limiting process: The phosphatase activity in the perturbed module is not the slowest. 2) Constraints on the initial conditions: The kinase activity needs to be fast enough such that each module is saturated even with fast phosphatase activity and upstream information is attenuated. We discussed the relevance of such robustness to several biological examples and the validity of the above conditions therein. Given the applicability of dynamics robustness to a variety of systems, it will provide a general basis for how biological systems function dynamically.

**Author Summary:** Cells use signaling pathways to transmit information received on its membrane to DNA,and many important cellular processes are tied to signaling networks. Past experiments have shown that cells’ internal signaling networks are sophisticated enough to process and encode temporal information such as the length of time a ligand is bound to a receptor. However, little research has been done to verify whether information encoded onto temporal profiles can be made robust. We examined mathematical models of linear signaling networks and found that the relaxation of the response to a transient stimuli can be made robust to certain parameter fluctuations. Robustness is a key concept in 1/15 biological systems it would be disastrous if a cell could not operate if there was as light change in its environment or physiology. Our research shows that such dynamics robustness does emerge in linear signaling cascades, and we outline the design principles needed to generate such robustness. We discovered that two conditions regarding the speed of the internal chemical reactions and concentration levels are needed to generate dynamics robustness.

## Introduction

Robustness is one of the most important concepts in biological systems. In general, it is the ability of an organism to maintain a state or behavior against external or internal perturbations, and many frameworks of robustness have emerged [1–7]. Homeostasis, for example, is the ability of an organism or a cell to maintain a certain state, such as its body temperature or calcium content, against external environmental changes. In fact, numerous mechanisms have been uncovered that are adopted to regulate its internal environment against external perturbations. In developmental biology, differentiated cellular states are known to be robust to disturbances, as was pioneered in the study by Waddington, who described the cell differentiation process as a ball rolling down an epigenetic landscape to settle into a stable valley [8]. This is a metaphorical representation of robustness often used, while in terms of dynamical systems theory, one mathematical formulation for static robustness can be described as an orbit being pulled into a stable attractor. The robustness discussed therein concerns the stationary state, and thus is regarded as *static robustness.*

In biology, however, both the static cellular states and dynamic processes are important to make certain responses against external changes robust and to ensure proper development. Waddington coined the term homeorhesis for such dynamics robustness for a transient time course [9]. Indeed, in the developmental process,temporal ordering of cell differentiations and their timing are robust. Besides the developmental process, cellular responses against external stimuli are often robust to perturbations since these time courses are often relevant to cellular function. Despite 22 the importance, such *dynamics robustness,* i.e., robustness in the temporal course, is little understood as compared with extensive studies on static robustness. Here we study dynamics robustness, the insensitivity of transients to initial conditions or parameters. We adopt the term *dynamics robustness* as opposed to dynamic or dynamical robustness since those terms have been defined elsewhere in a different context. For example in [10], dynamic robustness refers to the insensitivity of a steady-state against changes in protein concentrations to distinguish from the robustness of a steady-state against gene deletions. We stress that our focus is on the 30 robustness of the dynamics themselves against parameter perturbations.

As a specific example for such robustness, we focus on signaling pathways of covalent modification cycles. Indeed robustness therein has been extensively investigated, as given by a recent review by Bluthgen and Legewie [11]. Although their review is focused on static robustness in signal transduction pathways, they also note that ideas of robustness with regards to generating an invariant temporal profile (dynamics robustness) has to be developed [11]. In fact, there are several experiments suggesting robustness in the transient properties of certain biochemical networks: Different transient profiles of input stimuli can elicit different phenotypic responses. For example, it was shown that the duration of activation could lead to two different responses in PC12 cells; transient activation leads to proliferation, and sustained activation leads to differentiation [12, 13]. In a similar manner, the duration that a MAPK cascade is stimulated can lead to different responses in yeast [14]. Moreover, temporal profiles of the p53 pathway, which is inactivated in almost all human cancer cells, are also reported to be drastically altered by the types of stresses administered to the cells and cause different responses depending on the dynamic profiles [15]. All of these experimental 46 reports suggest the need for studies on dynamics robustness. Beyond such experimental results, research in the last decade has shown that signaling cascades can theoretically encode information into their dynamic profiles and process such information as well [16, 17]. For these dynamical processes to function, the time courses need to achieve 50 a certain level of robustness.

By investigating a class of signaling cascade systems, we also propose a novel concept, *duration robustness,* as a quantitative manifestation of dynamics robustness, wherein the duration of a response upon inputs is robust against perturbations. In a general class of cascading systems, we showed that duration robustness is an emergent property: Downstream modules are shielded from perturbations in the kinase activity in the upstream layers. Here, the organization of the fast and slow kinetics resulting in a rate-limiting module, is primarily responsible for such robustness. In a linear signaling system, by having fast kinase activity, the output time courses were shown to be robust to perturbations in the phosphatase activity. We uncovered two necessary conditions for dynamics robustness and demonstrated that it can be observed in general linear signaling systems via protein modifications. Furthermore, we verified that dynamics robustness is a property of the well-known model of a MAPK network described by 63 Huang and Ferrell [18].

## Results

Our results are organized as follows: We study a simple model of basic linear signaling 66 cascade and see how perturbing the parameters in the model affect the relaxation time courses. We first focus on perturbing the phosphatase parameters as they are known to 68 tend to control the duration in a signaling cascade. We then show how perturbing the kinase activities affect the results. Next, we analyze the linearized and normalized model of the aforementioned basic linear cascade to determine what underlying features of the cascade architecture causes dynamics robustness. From this analysis, we derive the conditions under which dynamics robustness is expected to emerge. Finally, we examine a more complicated mass-action model of a MAPK cascade to verify whether the results observed in the simple model are indeed features of a more biologically inspired model.

## Dynamics Robustness in the Heinrich Model

We first examined the Heinrich model of a general, linear signaling cascade (a detailed description can be found in the Methods Section). The basic idea is that a stimulus, the concentration of *E*_0_, activates a kinase, i.e., converts *M*_0_ to 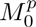, which goes on to activate a kinase downstream. This process occurs in three steps, and the concentration of the final activated kinase, 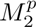, is considered the output response.

When time *t<* 0, a constant input 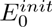 is applied to the system until 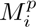 at each layer reaches the steady-state concentration, which we define as 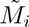. At time *t* = 0, *E*_0_ is set equal to zero and the system begins to relax into a deactivated state. Because it has been reported that phosphatase activity controls the duration more than the kinase activity [19, 22], we individually perturbed the total phosphatase activity at each layer and computed the new temporal profile to see if it remains robust. The parameters were chosen to reflect the same organization as the biologically relevant MAPK cascade parameters reported in [18] (see Supporting Information); the kinase activities are relatively fast, and the phosphatase rate constants are organized relatively as fast-slow-fast in the three stage setup. The specific *β* values from this parameter set correspond to the black circles in Figs. 1D-F. For clarity, the Heinrich model parameter definitions are given in Table 1.

**Table 1.**
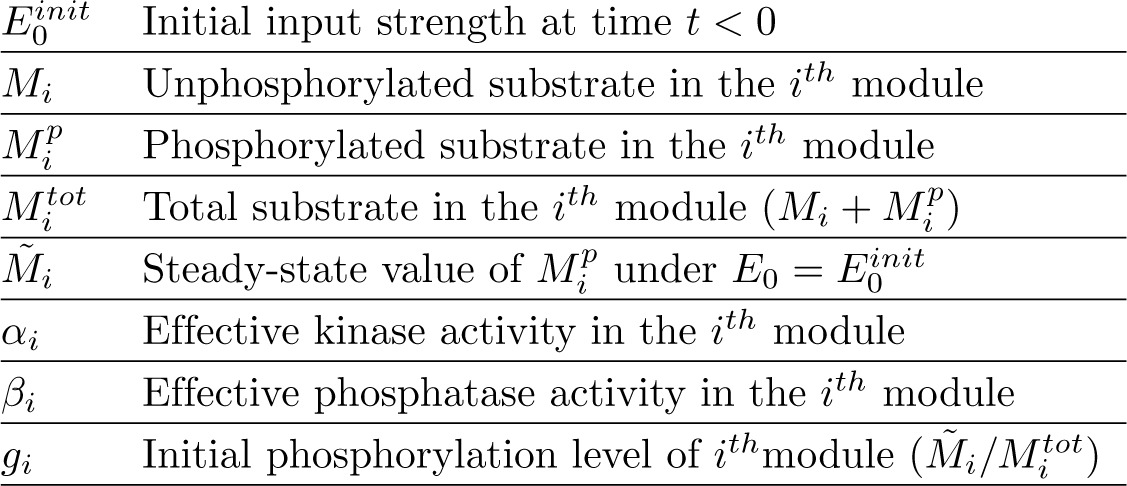
Parameters in the Heinrich model.

**Figure 1.**
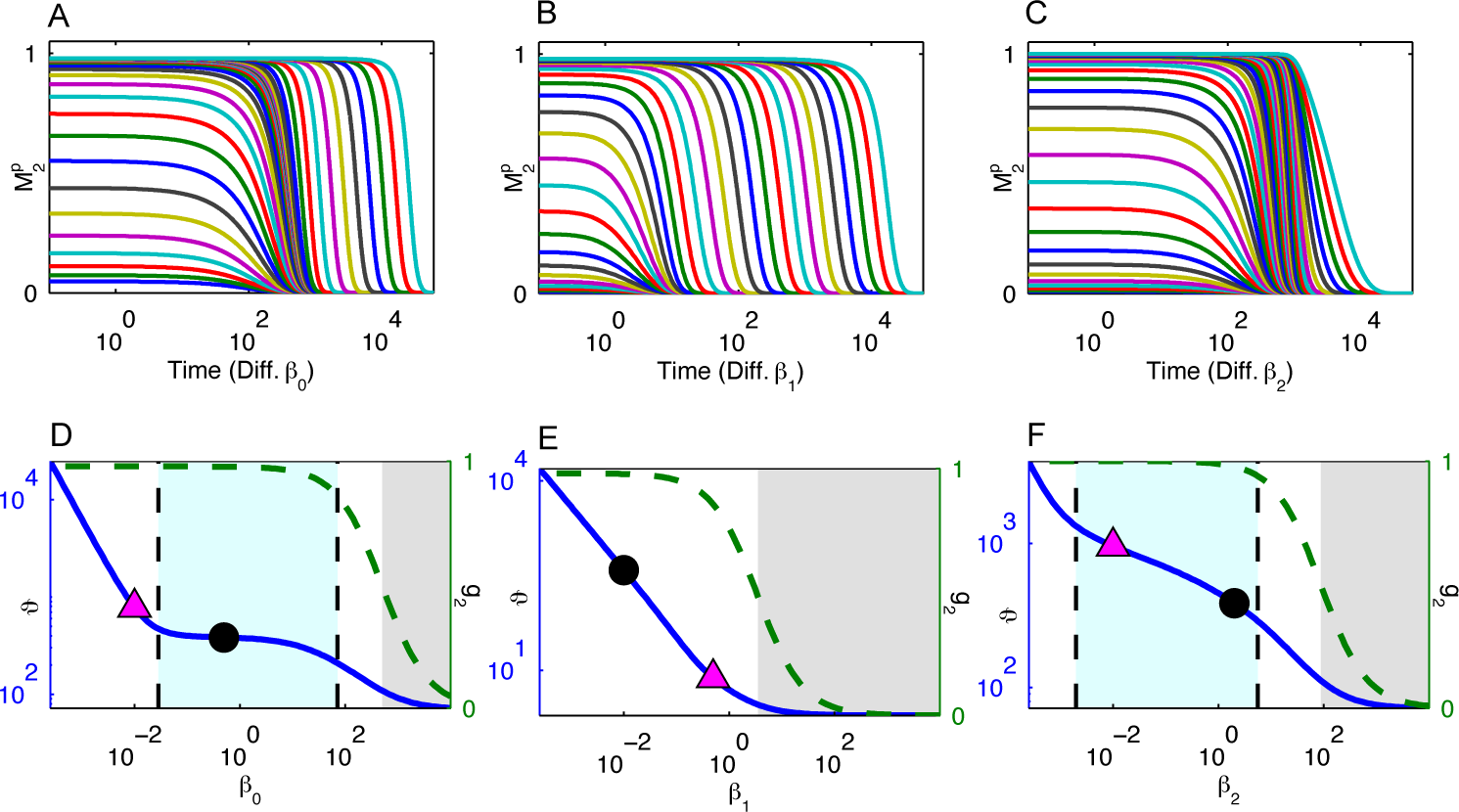
Dynamics Robustness in the Heinrich model. (A, B, C) The temporal profiles of the response relaxation for different values of *β*_0_ (A), *β*_1_ (B), and *β*_2_ (C). Different colors indicate the time courses for different *β_i_* values from the set 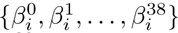 = {10^40^,10^38^,…, 10^−36^} where 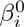 is for the lower leftmost line, and 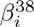 is for the top rightmost line. (D, E, F) The duration, *ϑ*, as a function of *β*_0_ (D), *β*_1_(E), and *β*_2_ (F). *ϑ* is plotted for the left *y*-axis. The initial phosphorylation level of *M*_2_ (i.e., *g*_2_) as a function of *β_i_* corresponds to the right *y*-axis. When *g*_2_ < 0.5, we consider the system to be in a deactivated state and color this region in gray. In between the vertical dashed lines, the system is in an activated state and the magnitude of the logarithmic gain of *ϑ* with respect to *β_i_* is less than 0.3. A magenta triangle indicates where *β_i_* becomes the minimum *β* value. A black dot indicates the *ϑ* with the parameters from Table S2.

The results for the Heinrich model are plotted in Fig. 1. There is an interesting parameter region where the temporal profiles are close together despite *β*_0_ decreasing from 10^2^ to 10^−2^ (Fig. 1A). There is a similar parameter region for *β*_2_ (Fig. 1C). In a certain range of *β*_0_ and *β*_2_, the temporal profiles do not change based on the phosphatase activity. See Fig S2 for the difference of the temporal profiles against changes in the phosphatase activity. However, there is no such parameter region in which the temporal profiles are not changed when *β*_1_ is perturbed (Fig. 1B). The temporal profiles of the output, 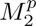, show dynamics robustness against changes in the phosphatase activity in the first and third layers, i.e., the time course profiles are robust to perturbations in *β*_0_ and *β*_2_.

As a simpler, analytically tractable measure for the robustness of the temporal profiles, we numerically computed the half-life, *ϑ*, which, as in [19], is defined to be the duration of the response. We plotted *ϑ* as a function of *β_i_* on a log-log scale in Figs. 1D-F. In this paper, we focus our discussion on the relaxation process of strongly activated cascades because the dynamics of a weakly activated signaling cascade are fundamentally different, and do not involve a significant relaxation time course. To clearly describe the criterion of activation, we introduce the parameter 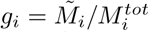, which is the ratio of the phosphorylated substrate to the total substrate in *i^th^* module. As we are interested in the response dynamics of the cascade, the initial activation *g*_2_ should be sufficiently high. Henceforth, we use the criterion that the cascade is activated if *g*_2_ > 0.5 (although the value 0.5 itself is not essential). We color the inactivated region in gray in Figs. 1D-F and focus on the dynamics in the region of strong activation.

As an indicator of robustness of *ϑ* against perturbations in *β_i_* we use the notion of logarithmic gain (see Methods section). In a typical case, such as in a single layered cascade, the logarithmic gain of the duration versus phosphatase activity would be −1, i.e., the duration is inversely proportional to the phosphatase activity, as expected by the relaxation form exp(−*βt*). The regions between the dashed vertical lines in Figs. 1D and F represent where the magnitude of the logarithmic gain is less than 0.3, which is distinctly smaller than 1. These flatter slopes indicate that the duration is 122 robust against changes in *β*_0_ and *β*_2_.

In Figs. 1D and F, the black circle, which represents the *β_i_* value from Table S2 and the corresponding *ϑ* value, is in the region of duration robustness, which means that with this parameter set reflecting actual kinetics in signaling cascades, the duration is robust to perturbations in the phosphatase activity in the first and last layer of the cascade (*β*_0_ and *β*_2_ respectively). However, the second module is sensitive to perturbations in the phosphatase activity.

In all three cases, there are common features in the plots of the duration. As mentioned earlier, if the phosphatase activity in any layer is too high, then the system is in an inactivated state, which is colored in gray in Figs. 1D-F. On the other hand, if the phosphatase activity in the *i^th^* layer is too low, then the logarithmic gain of the duration against *β_i_* is roughly −1, i.e., the duration of the response is strongly dependent on the rate-limiting module in the cascade. In Figs. 1D-F we plotted a magenta triangle at the value where *β_i_* becomes less than all other *β* values, and the logarithmic gain indeed becomes —1 near this point. However, the upper bound for the phosphatase concentration that exhibits duration robustness cannot be described by the rate-limiting effect only.

## Effects of the Kinase Activity on Duration Robustness

Although linear signaling cascades can show duration robustness against perturbations in the phosphatase concentrations, it is still unclear what effects the kinase activity has. Therefore, we computed the duration versus phosphatase activity (ϑ vs. *β_i_*) for different values of *α_i_* (Fig. 2).

**Figure 2.**
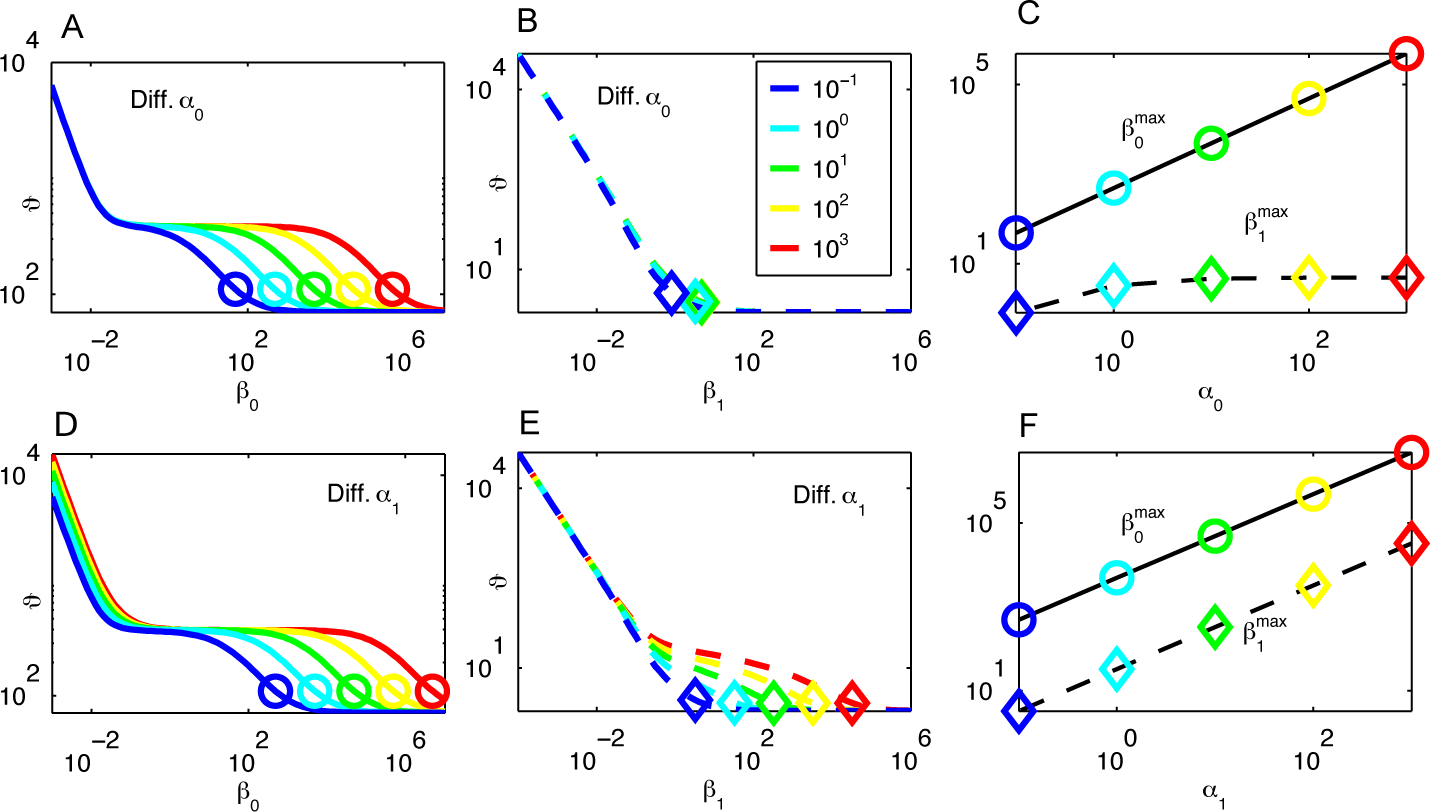
Effects of the Kinase Activity on Duration Robustness in the Heinrich Model. (A, B) Duration, *ϑ*, as a function of *β*_0_ (A) and *β*_1_ (B) with varied *α*_0_. Different lines indicate *ϑ* for different *α*_0_ values as given by the inset box in (B). Circles and diamonds represent 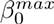 and 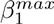, respectively. (C) 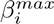 as a function of *β*_0_. A solid line and a dashed line are 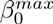 and 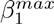, respectively. The circles and the diamonds correspond to these symbols in (A) and (B). (D, E) Duration, *ϑ*, as a function of *β*_0_ (A) and *β*_1_ (B) with varied *α*_1_. (F) 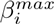 as a function of ai. Same colors, lines and symbols are adopted as (A), (B), and (C).

Increasing *α*_0_ expands the interval of duration robustness for *β*_0_, since the upper limit is increased while the lower limit remains fixed (Fig. 2A). This increase of *α*_0_, however, does not expand the duration for varied *β*_1_ (Fig. 2B). On the other hand,increasing ai expands the interval of duration robustness both for *β*_0_ and for *β*_1_: the slope of *ϑ* against *β*_1_ is flatter, resulting in the appearance of the region for duration robustness for *β*_1_.

Here, the upper limit of duration robustness is roughly given by the largest value of *β_i_*, which we call 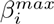, at which the system is activated. The 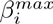 values are marked in Figs. 2A, B, D, and E. By using the criterion of *g*_2_, 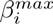 is given by the maximal value of *βi* that satisfies *g*_2_(*β*_1_) > 0.5. 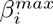 is then used as an indicator for the upper limit of the interval of duration robustness. To derive an expression for 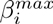, we see that *g_i_*, the phosphorylation level at each stage, can be written as a sequence of iterations:

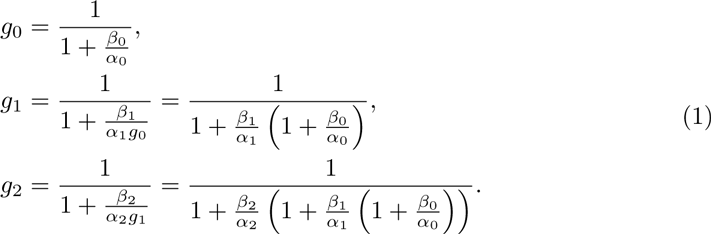

If changes in the kinase activity cause changes in 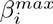 then the region of duration robustness will change as seen in Fig. 2. The iterative nature of Eq. 1 demonstrates how upstream information is shielded. It clearly shows that increasing *a_k_* will proportionally increase 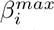 *only* for *i* ≤ *k*. For *i* > *k*, increasing *α_k_* has a negligible effect on 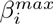 and hence, has a negligible effect on the interval of duration robustness for *β_i_*. This is somewhat counterintuitive because one usually considers alterations propagating downstream in a linear signaling cascade, whereas alterations in the kinase activity affects the range of duration robustness only in upstream modules. This is because the constraint for initial conditions back-propagates. In general, having fast kinase activity in the downstream modules is ideal if one wishes to generate a region of duration robustness against the upstream phosphatase activities.

## Duration Robustness in a Linearized Model

To better understand how duration robustness is generated and the criteria needed, we analyzed the linearization of the Heinrich model about the origin, the only equilibrium point once the stimulus is removed. Duration robustness is also a property of the linear model as can be seen in Fig. 3. The global linearization of the Heinrich model is a significant departure, and the time-course profiles for the linear case are drastically different from the ones for the nonlinear Heinrich model. In particular, the conserved quantities in the nonlinear model are no longer conserved in the linear model. However,the plots of the duration against the phosphatase activity in Figs. 3 are remarkably similar to those in Figs. 1. (The plots of the time-courses are provided in the Supporting Information, Fig. S3). This strongly suggests that the nonlinear kinetics are not important for duration robustness, although we will show that the nonlinearity of *g*_2_ as a function of *β_i_* does play a crucial role.

**Figure 3.**
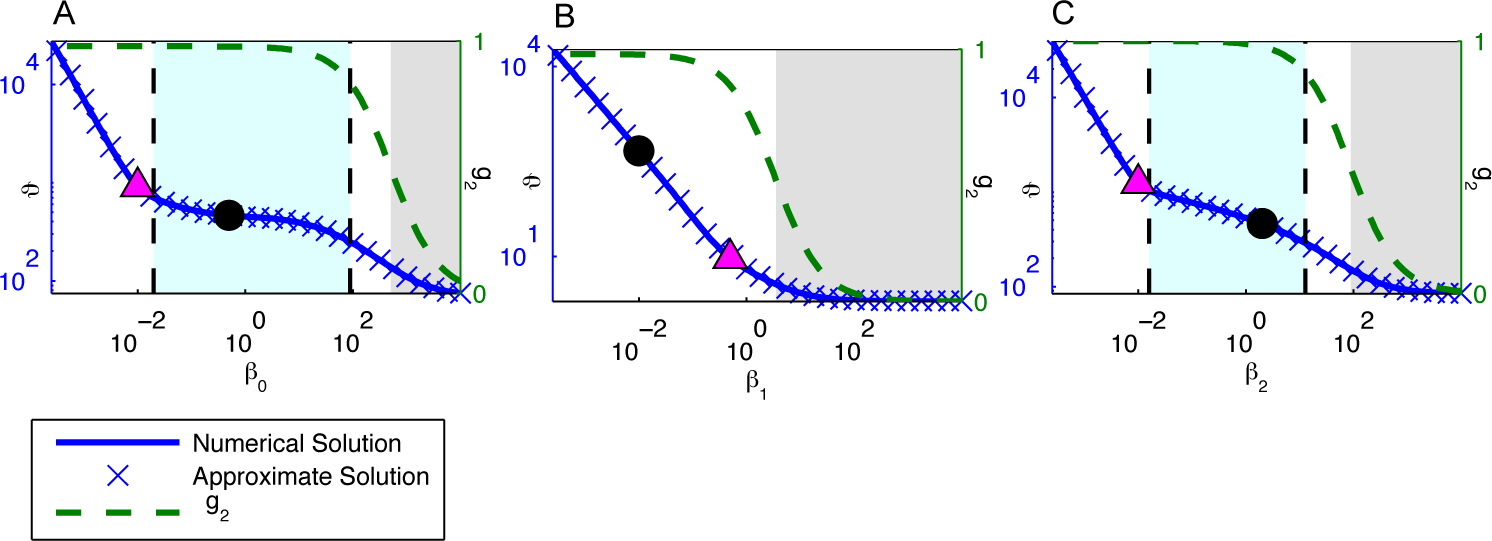
Duration Robustness in the Linearized Heinrich Model. The duration, *ϑ*, and the initial phosphorylation level, *g*_2_, as a function of *β*_0_ (A), *β*_1_ (B), and *β*_2_ (C). *ϑ* and *g*_2_ plotted in a similar manner of Fig.1. The approximation of the duration given in Eq. 2 is plotted as blue ×’s. Plots of the time course profiles are provided in the Supporting Information (Fig. S3).

If *α*_0_ ≠ *α*_1_ ≠ *α*_2_ the normalized solution 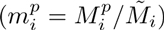 is just a linear combination of exponentials:

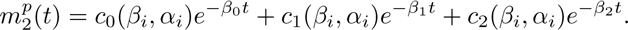

The duration (the time *ϑ* such that 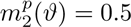) can be approximated by:

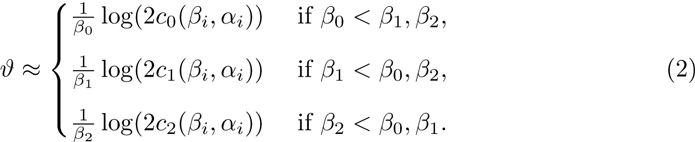

The pertinent question is how *ϑ* is made robust to changes in *β_i_*. If *β_i_* is the minimum *β* value, then the duration according to Eq. 2 is roughly inversely proportional to *β_i_*, which means that the logarithmic gain is going to be around −1. In fact, in the limit as *β_i_* goes to 0, the logarithmic gain converges to −1. In this case, accordingly, there is no duration robustness. Hence, to have duration robustness against *β*_1_, the first constraint is which we refer to as the *constraint on the rate-limiting process.*

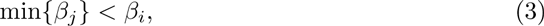

The lower limit of the *β_i_* interval in which duration robustness emerges is determined by this rate-limiting condition; however, this condition is not sufficient to determine the upper limit of the interval of *β_i_*. As already discussed, the initial phosphorylation level *g*_2_ at the output layer has to be sufficiently activated, and as shown in Fig. 3, the upper limit is strongly related to this initial phosphorylation level *g*_2_. Indeed, we can use Eq. 2 to understand this behavior analytically. Suppose that *β_k_* is the minimum *β* value and that *i* ≠ *k.* Then the logarithmic gain is given by:

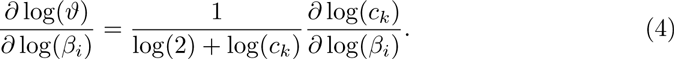

Therefore, if the logarithmic gain of *c_k_* with respect to *β_i_* is small, then duration robustness will emerge. As shown in the Supporting Information,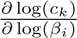 is strongly dependent on — 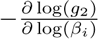.

If *g*_2_ has a sigmoidal nature as seen in Fig. 3, then it has two regions where it is relatively constant with respect to *β_i_* and a transition state between the two relatively constant regions. If this transition occurs before the module becomes rate limiting, then duration robustness will emerge because *g*_2_ will have a weak dependence on *β_i_*. As mentioned previously in relationship with Eq. 1, changes in upstream kinase activity have a negligible effect on *g*_2_, i.e., upstream information is shielded. Hence to increase the transition point and expand the region of duration robustness in the *i^th^* module, it is necessary that there exists some *j* ≥ *i* such that *β_j_* ≪ *α_j_*. In other words, it is necessary that in a module downstream, the kinase activity relative to the phosphatase activity needs to be very fast. We refer to this constraint regarding *g*_2_ as the *constraint on initial conditions.*

The arguments based on Eqs. 1 and 2 can be extended to an N-stage cascade, and the conditions needed to generate duration robustness in the *i^th^* module can be summarized as where the first condition represents the constraint on the rate-limiting process, and the latter two conditions give the constraint on the initial conditions.

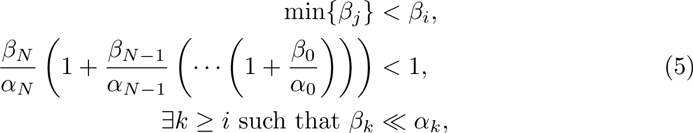

The arguments made for the linearization can also be extended to general linear signaling cascades. The rate-limiting condition can easily be understood using slow 206 manifold theory. The eigenmodes of a linear signaling cascade are proportional to the 207 phosphatase activity. Likewise, the phosphorylation levels at each stage do display a switch-like nature. Because the kinase activity controls the phosphorylation levels, both constraints, i.e., the rate-limiting condition and the constraint on the initial conditions, will also be necessary in any model of a linear signaling cascade.

Note that while both the original and linearized Heinrich models display duration robustness, the original Heinrich model displays a stronger type of dynamics robustness in the sense that the time-course profiles themselves are robust to changes in *β_i_* under certain conditions (see Fig. 1 and Fig. S3). This is mainly because in the linear model,the response is unsaturated and can vary, whereas the response in the nonlinear model is saturated, bounded, and decreasing for all relevant parameter regimes.

## Dynamics Robustness in the Huang Ferrell Model

To verify the general results on a more biologically inspired system, we next examined the Huang Ferrell (HF) model [18] of a linear signaling cascade (for a detailed description, see Supporting Information). This model is a complete mass action description of a MAPK signaling pathway, which is a linear cascade with three layers. The middle and last layers represent double phosphorylation events, which lead to ultrasensitivity [18]. The HF model also explicitly assumes that a phosphatase at each layer removes the active phosphate groups, and thus, the phosphatase activity is directly proportional to the total phosphatase concentration, 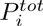, in each layer. The results in Fig. 4 demonstrate that duration robustness is also a property of the HF model. There are parameter regimes where the duration of the relaxation is insensitive to perturbations. Like the original Heinrich model, the HF model also displays dynamics robustness in which the time-course profiles themselves are robust to changes in the phosphatase concentrations. Figs. 4(C) and (F) show that the last layer in the HF model has slightly stronger dynamics robustness than the Heinrich model. This suggests that higher nonlinearities in signaling cascades may enhance dynamics robustness.

**Figure 4.**
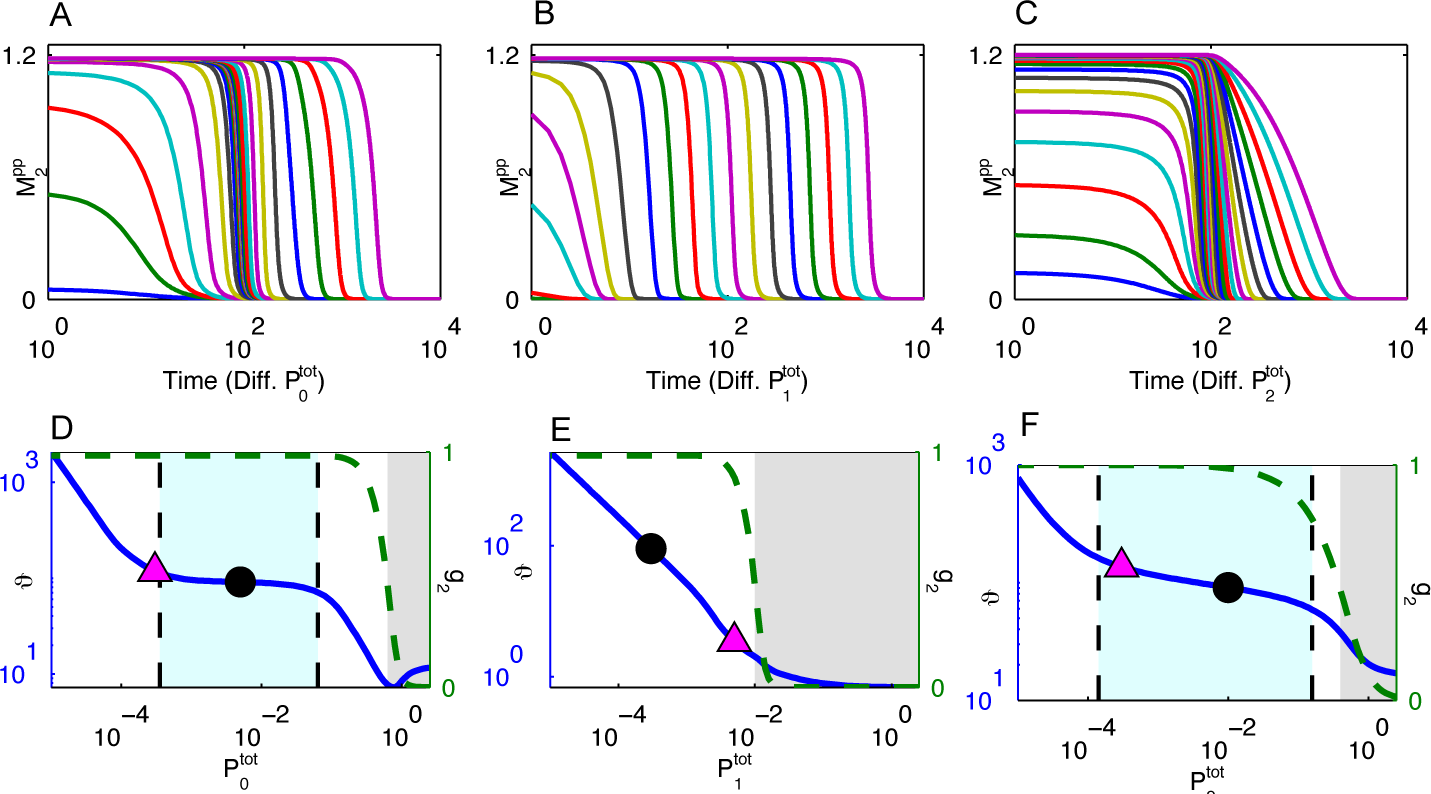
Dynamics Robustness in the Huang Ferrell Model. (A, B, C) The temporal profiles of the response relaxation for different values of 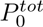, 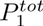, and 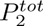 respectively. Different lines are for different values of 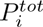 taken from the set {10^0.4^,10^0.2^,…10^−5^}. (D, E, F) The duration, *ϑ*, and the initial phosphorylation level, *g*_2_ are plotted as a function of 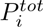, in the same way as Fig.1.

## Discussion

In the present paper, we have demonstrated that *dynamics robustness,* i.e., the insensitivity of the time courses against changes in certain parameters, is observed in the relaxation process of signaling cascades. By using a general linear cascading system, the time course of the output layer downstream is shown to be almost insensitive to changes in upstream parameters. As a consequence of dynamics robustness, the duration in which activated state lasts is also robust to parameter changes, a property we termed *duration robustness.* By analyzing the cascading process, the conditions for duration robustness are given by the constraint on the rate-limiting process and on the 242 initial conditions. Since multiple layers are needed to generate duration robustness, this suggests that this property is a byproduct of how temporal information is processed downstream.

## Conditions for duration robustness

We have shown that linear signaling cascades of varying complexity display duration robustness against perturbations in the phosphatase activity in the *i^th^* stage, and that two main conditions are responsible for this phenomenon:

## 1) The constraint on the rate-limiting process

The phosphatase activity in the *i^th^* stage, *β_i_*, should not be the minimum *β* value. This unfortunately means that the slowest module in a linear cascade will not display duration robustness. This constraint determines the lower limit for the range of duration robustness, i.e., *β* < *β*_*i*_. If *β*_*i*_ is the minimum *β* value in the cascade, then the duration time is inversely proportional to *β*_*i*_ as described by usual relaxation processes. In general linear signaling cascades, this means that the phosphatase activity in module *i* should not be the slowest. For certain parameter regions, our results are contrary to the idea that upstream phosphatase activity controls the duration of the system more than downstream in a strongly activated cascade [19].

## 2) The constraint on the initial conditions

To achieve duration robustness, the initial phosphorylation level of the output layer also has to be robust. For the Heinrich model, the initial phosphorylation level, *g_i_*, is given as a sequence of iterations as Eq. 1, and if the kinase activity in some layer is sufficiently high, *g_i_* will be robust against changes in the upstream phosphatase activity. In other words, information at upstream layers is shielded by the strong kinase activity. This constraint determines the upper limit for duration robustness.

Intuitively, if the kinase activity is low, the phosphatase activity should be low enough to allow the cascade to be active. How large *β_i_* can be is largely determined by the kinase activity, *α_i_*. A stronger kinase activity allows the phosphatase to be at a higher level and the system to remain in an activated state. Although too low kinase activity changes the initial phosphorylation level, too high kinase activity has little effect, due to the saturation of the phosphorylation. This determines the upper limit of *β_i_* for the region of duration robustness.

In general, linear signaling cascades do display such saturation, as a result of conservation of the substrate at each layer, and as for the Heinrich model, the steady-state phosphorylation level in *i^th^* layer could be given as

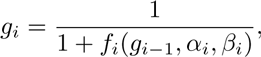

where, *f_i_*(*g*_*i*−1_, *a_i_, β_i_*) is a decreasing function of the kinase activity, *α_i_*, and an increasing function of the phosphatase activity, *β_i_*. In this case, increasing the kinase activity downstream can shield the upstream information, and then lead to duration robustness against changes in the phosphatase activity in upstream modules. This is interesting as changes appear to be propagated upstream. This type of downstream-to-upstream perturbation transference was reported in the steady-state 279 concentration of linear cascades as *retroactivity* [23], whereas our upstream propagation in the duration robustness is a different type of retroactivity since it concerns with the initial condition for shielding the information.

## Biological Relevance

Our results showed that within the range of a biologically relevant parameter set of a MAPK signaling pathway reported in [18], the duration and temporal profile of a strongly activated response are robust against perturbations in the phosphatase activities in the first and last modules. Past research has shown that temporal profiles of signaling cascades upon different inputs can lead to drastically different behaviors in cells. As mentioned earlier, transient versus sustained activation leads to different developmental responses [12, 13], and the behavior of transients in the p53 pathway is important to understanding certain types of cancer [15]. Our theory of dynamics robustness suggests that the transients involved in such decision processes can be robust to internal fluctuations in the concentrations of enzymes. We claim that stronger kinase activities are important for generating robust temporal profiles and that such a relationship will be verified experimentally.

The fast-slow-fast organization of the kinetics in the three-stage cascade is not necessary for dynamics robustness since it is observed in two-stage cascades as well. We looked at other kinetic organizations and their results intuitively agreed with the results 298 in this paper; the rate-limiting module tends to control the duration and the other modules display robustness under the constraints discussed. Whether the fast-slow-fast 300 organization is a byproduct of another selected property or is selected for a beneficial trait regarding dynamics robustness is unknown. However, one possible benefit is the emergence of a plateauing response as observed in Fig. 4(A) and Fig. 1(A). In this 303 plateauing behavior, the response remains in a quasi-steady state before decaying exponentially. It is possible that a three-stage linear cascade may be used to store information in one of these reliably timed plateaus. Dynamics robustness may explain the reliability of the response, but future work is needed to explain the mechanism of the plateauing response and its relationship with dynamics robustness. This type of plateauing response has been discussed before as kinetic memory in other biochemical 309 systems [24, 25] and such memory will also emerge in a linear cascade with a fast-slow-fast organization.

As a design principle, a signaling cascade with the conditions discussed previously are ideal for robust transients and this parameter organization is reflected in [18]. Since reliably timed transients are useful in signal processing, robustness would make such properties evolutionarily feasible. Indeed, a repetitive cascade structure would be easily evolvable by gene duplication [26, 27], wherein the function of the original cascade is safeguarded by robust parameters.

We focused on linear signaling cascades because of the recent interest in their temporal dynamics, but our idea of dynamics robustness can be generalized to any biochemical network. Two common properties of the cascade architectures we examined 320 were mass conservation and an active molecule working as a kinase downstream. This suggests that such robustness can be achieved by similar designs such as the two-component signaling network in bacteria, and may be a universal feature in biological signaling cascades via protein modifications. There are some biochemical processes which are known to be reliably timed, such as lysis of bacteria and chromosome segregation during mitosis [28, 29], and these reliably-timed processes might be considered as a demonstration of dynamics robustness. Such robustness should be an essential property to many biological systems, and the expansion of the present formulation will provide a future fruitful area of research.

The concept and explicit results of *dynamics robustness* we have presented here should be timely and of importance. In many biological phenomena, the time course, such as the response against external stimuli or the developmental process, is crucially important, and must be sufficiently robust to perturbations. This point has been noted before, but so far there is no theory for such dynamics robustness. For example, the scale invariance of time course has gathered much attention as fold-change-detection [30]. Dynamics robustness concerns with the insensitivity to external changes rather than the scale invariance of time courses, and does not require strong constraints as imposed in the fold-change-detection. Dynamics robustness can appear in a cascading system in general, by shielding upstream parameter changes. Thus it will have broader impacts and applications.

We demonstrated this dynamics robustness in standard models of signal transduction. As these models are based on experimental data, and agree rather well 342 with them, our dynamics robustness can be straightforwardly confirmed in cell-signaling experiments. Also considering the generality of our results, many other experimental topics will benefit from dynamics robustness.

## Models and Methods

We looked at different models of varying complexity. Although we use the nomenclature of kinases and phosphatases to represent the activating enzymes and deactivating enzymes, our model can be applied generally to any linear signaling cascade. We used mass action kinetics to simulate the chemical reactions, and all of our equations were solved using MATLAB’s (version R2009a) built-in numerical integrator ode15s.

### Heinrich Model

In the Heinrich model described in [19] and diagrammed in Fig. 5,the receptor, *E*_0_, converts *M*_0_ to 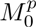, and 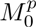 converts *M*_1_ to 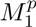, and 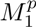 converts M_2_ to 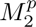, which is the output. The second order reaction rate at which *M_i_* is activated is 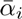, and the first order deactivation rate for 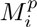 is *β_i_*. We assume that after an initial, constant stimulus and equilibration of the system, the receptor is immediately shut off and the system relaxes.

**Figure 5.**
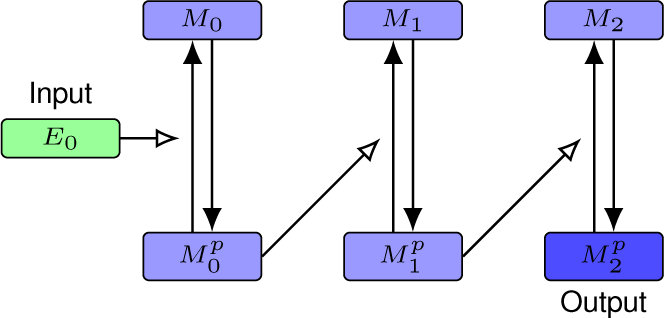
Diagram of the Heinrich Model. A linear signaling cascade is a biochemical network where the product of one reaction acts as an enzyme for a reaction downstream. The Heinrich model captures the basic essence of such an architecture. For time *t* < 0, the receptor, *E*_0_ receives a stimulus with strength 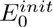. *E*_0_ then converts *M*_0_ to 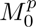 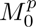 then converts *M*_1_ to 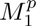, and 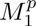 converts *M*_2_ to 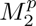. The concentration of 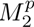 is considered the output response. After the system reaches a steady-state, at time *t* = 0, the stimulus is immediately removed, and the system then settles into a deactivated state.

There are a few major simplifying assumptions in this model that make it useful for examining the qualitative behavior of linear cascades. The assumptions are that the intermediate complexes formed by each kinase-substrate pair is negligible, that the backward reaction from the complexes is insignificant, and that the active phosphatase concentration is nearly constant. This means that the phosphatases and the intermediate complexes can be ignored, the desphosphorylation rate can be expressed as a first-order reaction rate, and that the sum of the inactive and active forms of each substrate is constant, i.e., 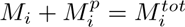 where 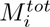 represents the total amount of substrate *M_i_*. Although these assumptions ignore some details, they enable us to analyze the models mathematically while still capturing the overall behavior of a signaling cascade. The corresponding set of equations post-stimulus is:

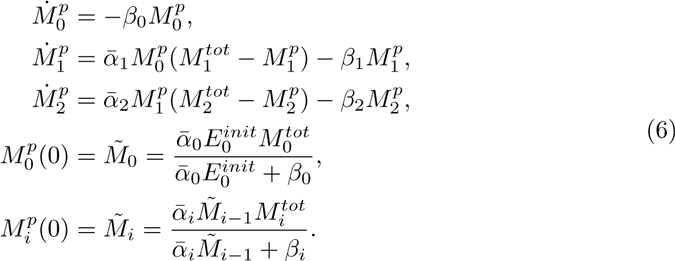

We note that Eqs. 6 is equivalent to Heinrich’s model, albeit with a slightly different form. An equivalent normalized model, i.e., where 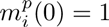, has the form:

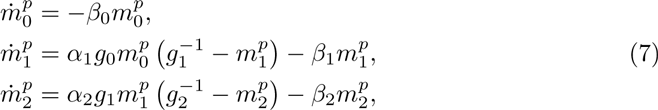

where 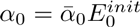 and 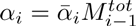 are the effective kinase activities.

### Logarithmic Gain

How robust a system is to a perturbation in a parameter is quantitatively measured by logarithmic gain. If one plots the dependent variable (say *y*) against a parameter (say *x*) on a log-log scale, then the logarithmic gain at a point is the slope of the tangent at that point. In other words, the logarithmic gain at a point *x*_0_ is 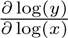 at *x*_0_. If *y* is inversely proportional to *x*, then the logarithmic gain will be −1. This concept has been used in systems biology to measure the robustness of steady-state concentration levels and transition times [20, 21], but here we use it to measure how robust the half-life of a linear signaling cascade is against parameter changes, which, to the best of our knowledge has not been done before.

## Supporting Information

### S1 Appendix

#### Additional Information

We provide a table of the parameters used in our simulations, additional figures, and equation derivations.

## Supporting Information

### 1 Huang and Ferrell Model

**Figure S1:**
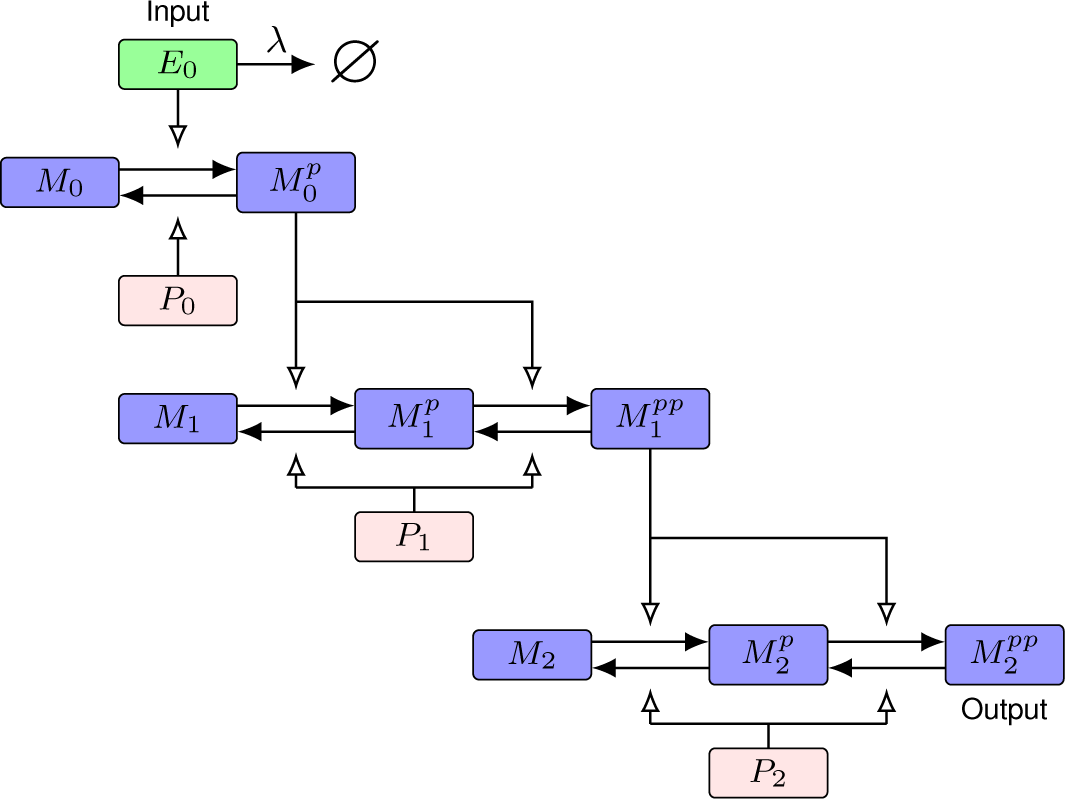
Diagram of the Huang and Ferrell Model. Unlike the Heinrich model, the HF model assumes double phosphorylation events, i.e., two phosphate groups are needed to fully activate *M*_1_ and *M*_2_. Once the system reaches an equilibrium state, the stimulus at the top layer is removed at a rate of λ.

In the original Huang and Ferrell (HF) model, an input stimuli, *E*_0_, activates a MAP-kinase at the top layer. We labeled this substrate, *M*_0_. We note that our labels are different from the labels Huang and Ferrell used so we could be consistent with other models in this paper. The activated form, 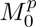, can be deactivated by *P*_0_, or it can go on to activate a MAP-kinase downstream, *M*_1_. Using *Xenopus* oocyte extracts as a model, Huang and Ferrell assumed that two phosphorylation events are needed to activate *M*_1_ and the next MAP-kinase downstream, *M*_2_. Likewise, the activated form,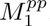, is deactivated by the phosphatase *P*_1_ in a two step process, and 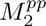 is dephosphorylated by *P*_2_ in a two step process. The 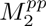 concentration is regarded as the response. The rate equations are derived by mass action assumptions. The kinetic parameters that Huang and Ferrell used, which reflect actual parameters experimentally derived, are listed in Table S1. To best illustrate the concept of dynamics robustness, the basal phosphatase concentrations we used are slightly different from the original Huang and Ferrell parameters.

As in the Heinrich model, we gave the system an initial input with strength 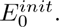 We allowed the system to reach an equilibrium state, and then removed the stimulus at a rate of λ. Because the HF model assumes complex formation, it was necessary to specifically remove the stimulus instead of simply setting it equal to zero as was done in the Heinrich model.

The stoichiometry of the Huang-Ferrell model [1] with enzyme destruction is given by:

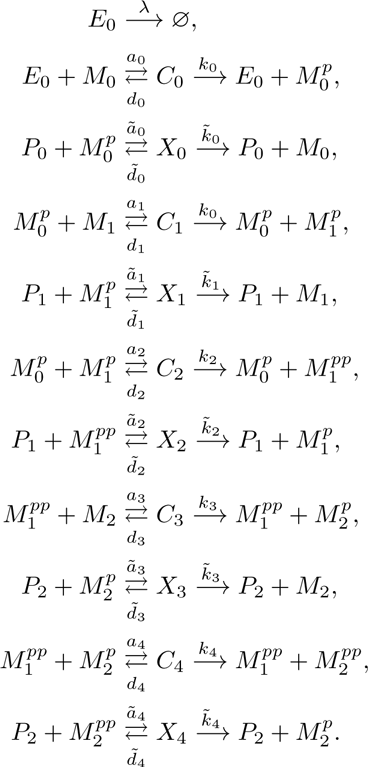

and the corresponding mass-action system is given by:

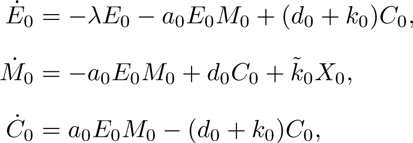

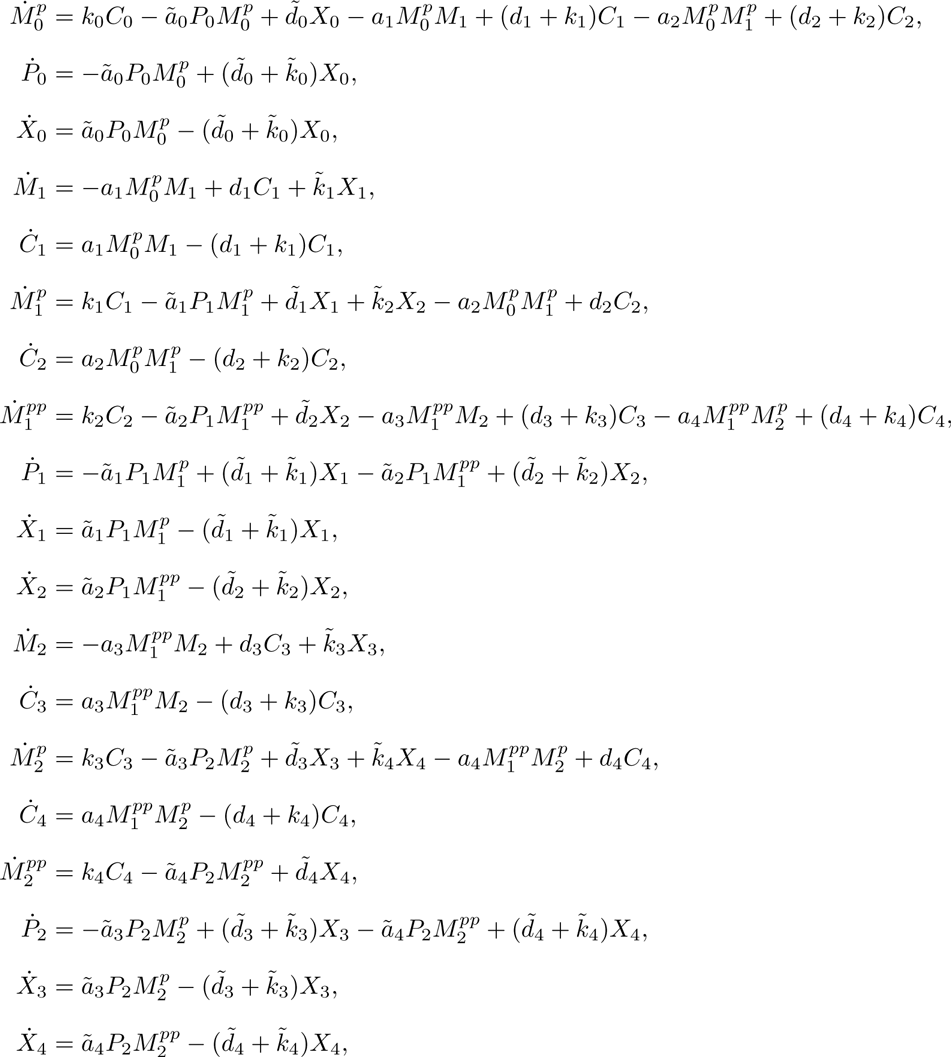

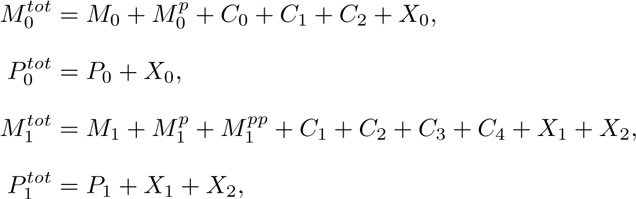

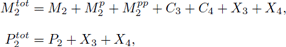

where the last six equations represent the conserved quantities.

### 2 Heinrich and HF Model Parameters

**Table S1.** Parameters used for the Huang and Ferrell model. The original parameters from [1] are displayed in the third column. The parameters used for our results are within a reasonable range of the original parameter set, but are slightly different to better exemplify dynamics robustness.

| Parameter | Units | Value From [1] | Base Value for Our Results |
| --- | --- | --- | --- |
| $E_0^{init}$ | $\mu M$ | — | $3 \times 10^{-3}$ |
| $M_0^{tot}$ | $\mu M$ | $3 \times 10^{-3}$ | $3 \times 10^{-3}$ |
| $P_0^{tot}$ | $\mu M$ | $3 \times 10^{-4}$ | $5 \times 10^{-3}$ |
| $M_1^{tot}$ | $\mu M$ | 1.2 | 1.2 |
| $P_1^{tot}$ | $\mu M$ | $3 \times 10^{-4}$ | $3 \times 10^{-4}$ |
| $M_2^{tot}$ | $\mu M$ | 1.2 | 1.2 |
| $P_2^{tot}$ | $\mu M$ | 0.12 | 0.01 |
| $\lambda$ | $(\text{min})^{-1}$ | — | 0, 100 |
| $a_i, \tilde{a}_i$ | $(\mu M \cdot \text{min})^{-1}$ | 1000 | 1000 |
| $d_i, \tilde{d}_i$ | $(\text{min})^{-1}$ | 150 | 150 |
| $k_i, \tilde{k}_i$ | $(\text{min})^{-1}$ | 150 | 150 |

**Table S2.**
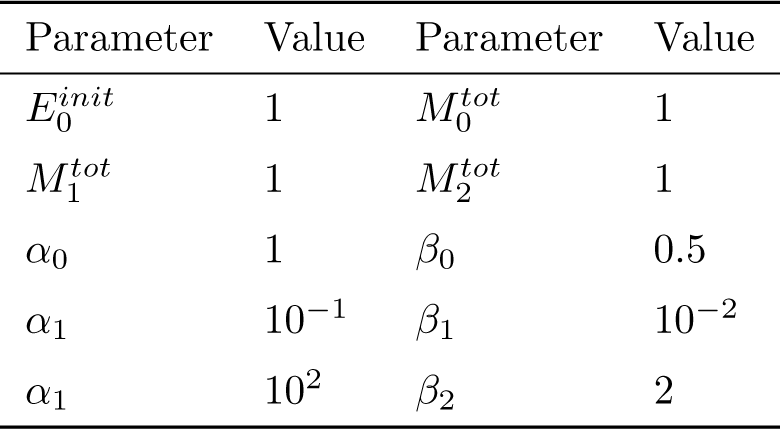
Parameters used for the Heinrich model [2]. These parameters were chosen to reflect the same organization of the kinetics from the Huang and Ferrell model.

| Parameter | Value | Parameter | Value |
| --- | --- | --- | --- |
| $E_0^{init}$ | 1 | $M_0^{tot}$ | 1 |
| $M_1^{tot}$ | 1 | $M_2^{tot}$ | 1 |
| $\alpha_0$ | 1 | $\beta_0$ | 0.5 |
| $\alpha_1$ | $10^{-1}$ | $\beta_1$ | $10^{-2}$ |
| $\alpha_1$ | $10^2$ | $\beta_2$ | 2 |

### 3 Complete Results for the Heinrich, Linearized, and Huang-Ferrell Model

Here we display the time-course profiles, the *L*^2^ norm between consecutive temporal profiles, the duration, and the logarithmic gain of the duration with respect to changes in the phosphatase activity for the Heinrich, Linearized Heinrich, and the HF models.

#### Heinrich Model

In Fig. S2(A), we set *β*_0_ and *β*_2_ equal to their base values from Table S2. We then integrated the Heinrich model for each of the *β*_0_ values taken from the set 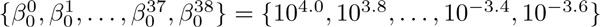. There is an interesting point where changing *β*_0_ does not alter the temporal profiles by very much. To measure how close the time-course profiles are, in Fig. S2(B), we computed the *L*^2^ norm between consecutive temporal profiles from Fig. S2(A). In other words, we computed

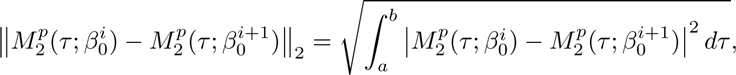

where τ = log_10_(*t*) and the integral is approximated using Matlab’s **trapz** method over the interval [−2, 4 + log_10_(6)].

In Fig. S2(C), we computed the half-life for the Heinrich model for each of the *β*_0_ values taken from the set 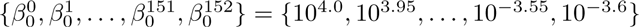.In Fig. S2(D), we numerically approximated the logarithmic gain of the duration versus changes in *β*_0_ using Matlab’s **diff** method on the results in Fig. S2(C).In Fig. S2(E-H), we repeated the experiment except setting *β*_0_ and *β*_2_ to their base values from Table S2 and varied *β*_1_. In Fig. S2(I-L), *β*_0_ and *β*_1_ are set to their base values from Table S2 and *β*_2_ is varied. The results show that there is an interval of P0 where the system is in an activated state and the temporal profiles and their duration are very robust against perturbations in *β*_0_. Likewise for *β*_2_, albeit with a weaker robustness compared to *β*_0_. However, no such interval exists for *β*_1_.

#### Linearized Heinrich Model

The same experiment as for the Heinrich model was performed for its linearization about the origin and the results are displayed in Fig. S3. Unlike in the Heinrich model, the temporal profiles are not very robust to changes in *β_i_* when the system is in an activated state. However, the results for the duration in Fig. S3(C,G,K) are remarkably similar to the results for the nonlinear model in Fig. S2(C,G,K).This suggests that while the nonlinear kinetics are important for achieving dynamics robustness in the Heinrich model, the nonlinear kinetics are not important for achieving duration robustness.

#### HF Model

The same experiment as for the Heinrich model was performed for the HF model and the results are displayed in Fig. S4. In Fig. S4(A), we set 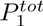 and 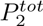 equal to their base values from Table S1. We then integrated the HF model for each of the 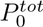 values taken from the set 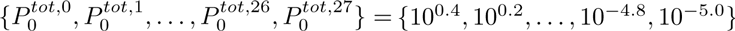. In Fig. S4(B), we computed the *L*^2^ norm in the difference between consecutive temporal profiles. We integrated over the log of time and over the interval [−2, 4]. In Fig. S4(C), we computed the half-life for the HF model for each of the 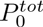 values taken from the set 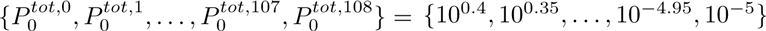. In Fig. S4(D), we numerically approximated the logarithmic gain of the duration versus changes in 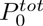. We then repeated the experiment in the same manner as in the Heinrich results to generate the other plots. The results are qualitatively similar as the Heinrich results in Fig. S2, which suggests that dynamics robustness and duration robustness are intrinsic properties of linear signaling cascades.

**Figure S2:**
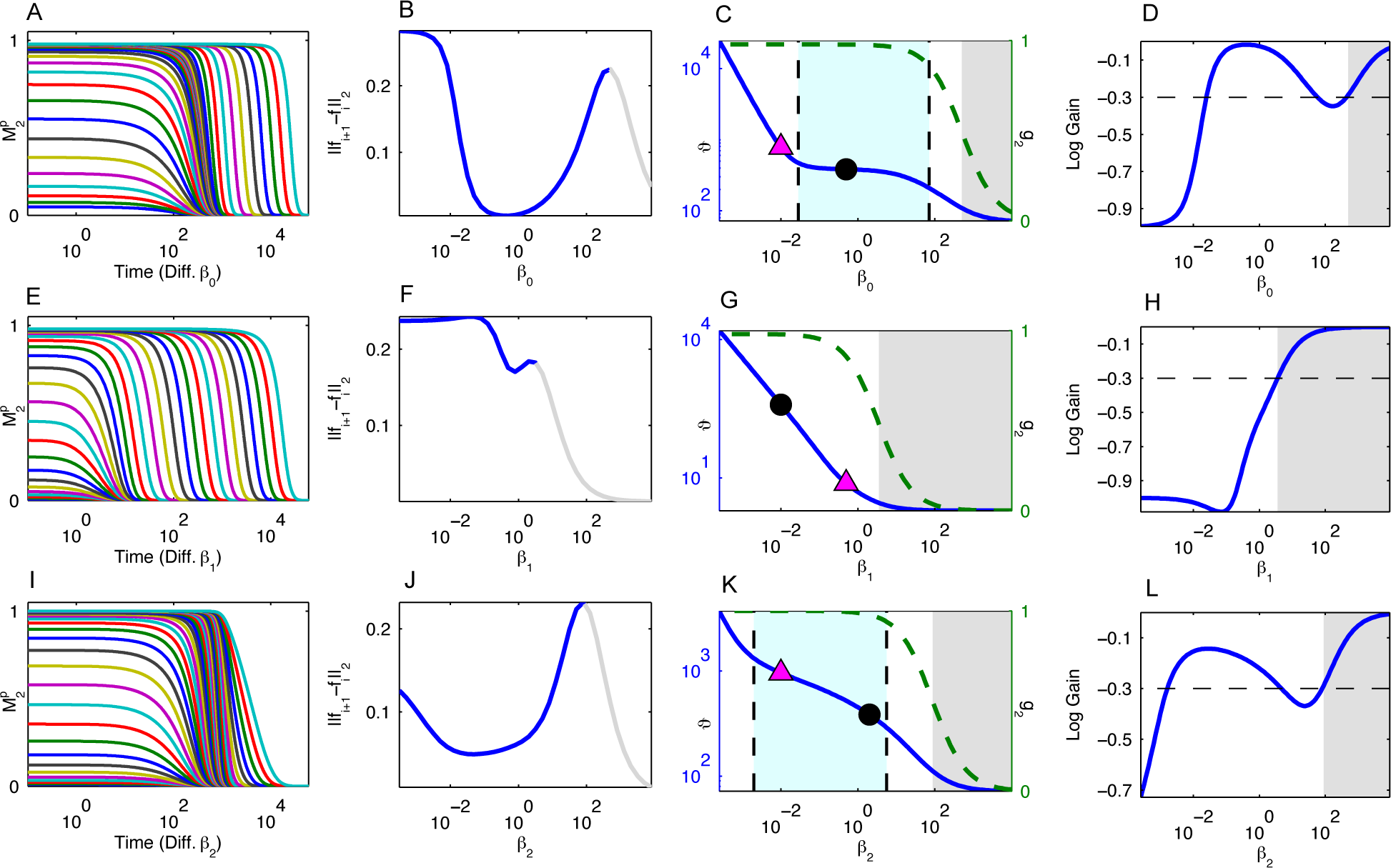
Complete Results for the Heinrich Model. (A, E, I) The temporal profiles of the response relaxation for different values of *β*_0_ (A), *β*_1_ (E), and *β*_2_ (I). (B, F, J) The consecutive similarity in the temporal profiles for *β*_0_ (B), *β_1_* (F), and *β*_2_ (J). The gray portion of the lines in (B, F, J) indicate that the system is in a deactivated state (i.e., *g*_2_ < 0.5) for those values of *β*_*i*_. (C, G, K) The half-life of the response as a function of *β*_0_(C), *β*_1_ (G), and *β*_2_ (K). The magenta triangle triangle indicates when the *β_i_* value becomes the minimum *β* value. The black dot represents the base *β_i_* value from Table S2. The grayed out region indicates that the system is in a deactivated state for those values of *β_i_*. The region between the dashed vertical lines indicate that the magnitude logarithmic gain of the duration against *β*_i_ is less than 0.3. (D, H, L) The logarithmic gain of the duration against *β*_0_ (D), *β*_1_ (H), and *β*_2_ (L).

**Figure S3.**
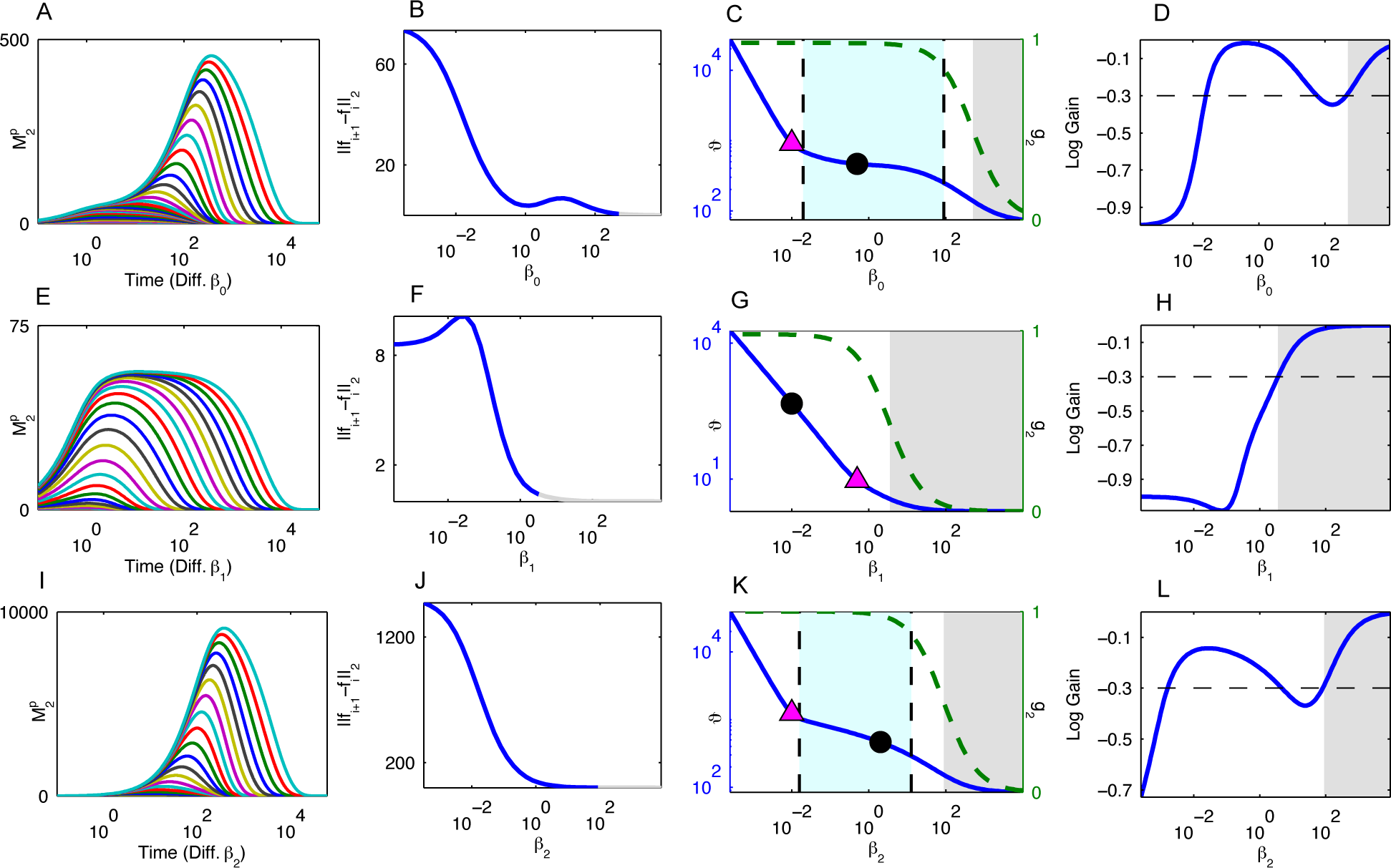
Complete Results for the Linearized Heinrich Model. (A, E, I) The temporal profiles of the response relaxation for different values of *β*_0_ (A), *β*_1_ (E), and *β*_2_ (I)- (B, F, J) The consecutive similarity in the temporal profiles for *β*_0_ (B), *β*_1_ (F), and *β*_2_ (J). The gray portion of the lines in (B, F, J) indicate that the system is in a deactivated state (i.e., *g*_2_ < 0.5) for those values of *β*_1_ (C, G, K) The half-life of the response as a function of *β*_0_ (C), *β*_1_ (G), and *β*_2_ (K). The magenta triangle triangle indicates when the *β_i_* value becomes the minimum *β* value. The black dot represents the base *β_i_* value from Table S2. The grayed out region indicates that the system is in a deactivated state for those values of *β_i_*. The region between the dashed vertical lines indicate that the magnitude logarithmic gain of the duration against *β_i_* is less than 0.3. (D, H, L) The logarithmic gain of the duration against *β*_0_ (D), *β*_1_ (H), and *β*_2_ (L).

**Figure S4.**
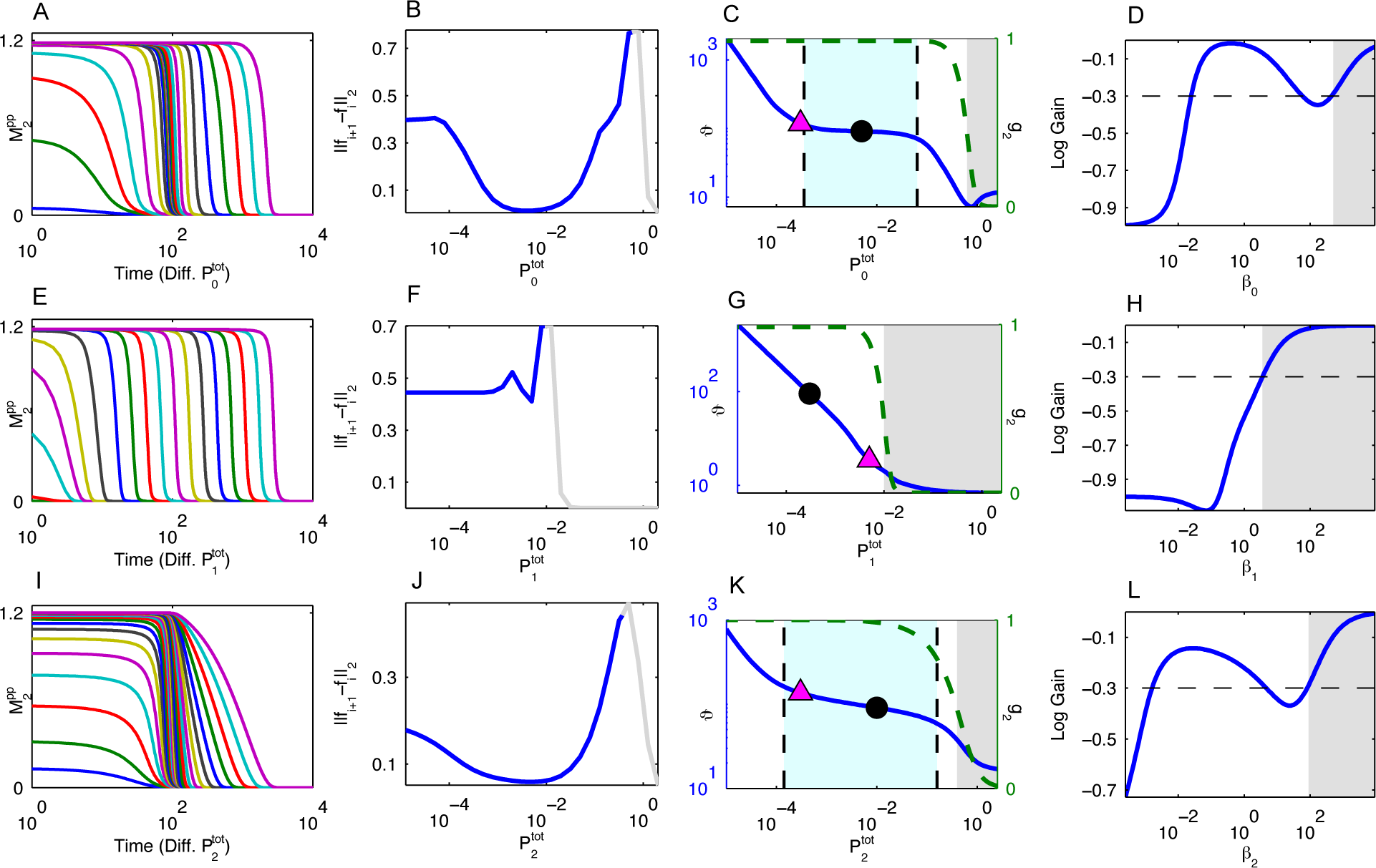
Complete Results for the HF Model. (A, E, I) The temporal profiles of the response relaxation for different values of 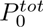 (A), 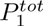 (E), and 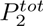 (I)- (B, F, J) The consecutive similarity in the temporal profiles for 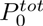 (B), 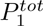 (F), and 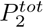 (J). The gray portion of the lines in (B, F, J) indicate that the system is in a deactivated state (i.e., *g*_2_ < 0.5) for those values of 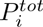. (C, G, K) The half-life of the response as a function of 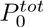 (C), 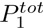 (G), and 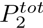 (K). The magenta triangle triangle indicates when the 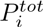 value becomes the minimum *P^tot^* value. The black dot represents the base 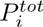 value from Table S1. The grayed out region indicates that the system is in a deactivated state for those values of 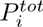. The region between the dashed vertical lines indicate that the magnitude logarithmic gain of the duration against 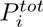 is less than 0.3. (D, H, L) The logarithmic gain of the duration against 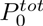 (D), 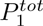 (H), and 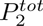 (L).

### 4 Derivation of Equations

#### 4.1 The largest value of *β_i_* at which the system is activated

The equations for 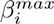 can be easily derived from the *g_i_* functions, and they also demonstrate how the kinase activities only alter the 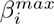 values and the corresponding regions of duration robustness upstream.

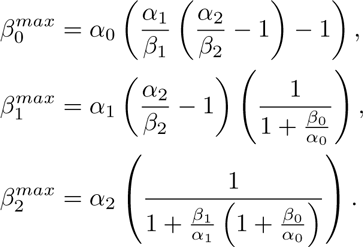

#### 4.2 Logarithmic Gains

In the main text it was shown that

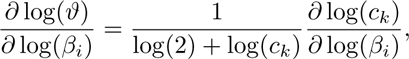

where *i* and *k* are such that *β_k_* is the minimum *β* value and *i* ≠ *k*. Our goal is to determine under what conditions is 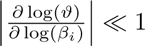

##### Case I: *β*_0_ is minimum *β* value, and duration robustness with respect to *β_i_*

First, let us consider the case when *β*_0_ is the minimum *β* value, and under what conditions will the duration be robust to changes in *β*_1_. We have that

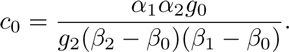

Hence,

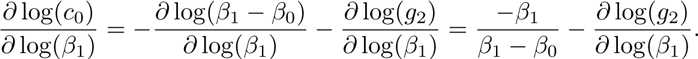

Therefore,

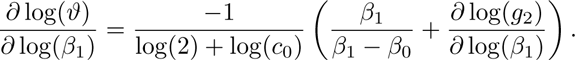

Thus, to minimize 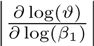 when *β*_1_ is sufficiently larger than *β*_0_,it is necessary to maximize *c*_0_ and to minimize 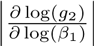. Now,

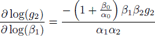

Hence, if

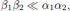

then 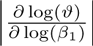 will be minimized.

##### Case II: *β*_0_ **is minimum** *β* **value, and duration robustness with respect to** *β*_2_

A similar argument as above shows that the condition needed is

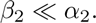

##### Case III: *β*_1_ is minimum *β* value, and duration robustness with respect to *β*_0_

In this case, we have that

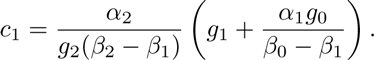

Hence,

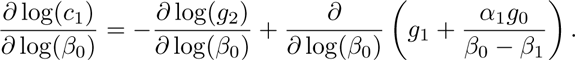

It can be shown that the second term is equal to

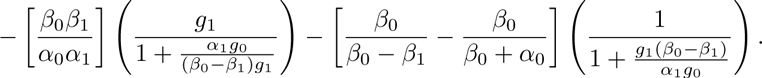

The terms in the parentheses are all less than 1. Hence, |log(2) + log(*c*_1_)| can be maximized and 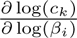 can be minimized if

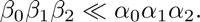

**Case IV**: *β*_1_ is minimum *β* value, and duration robustness with respect to *β*_2_. We have that

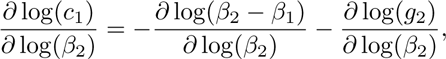

and the same argument as Case I can be used to show the condition needed is

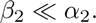

**Case V and VI**: *β*_2_ is minimum *β* value, and duration robustness with respect to *β*_0_, *β*_1_ We have that

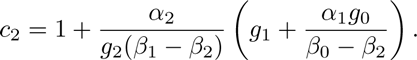

A similar analysis as above shows that the condition needed is that

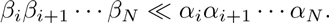

Summary In all, given the constraint that the cascade is considered activated, the constraint on the initial conditions can be grouped together as:

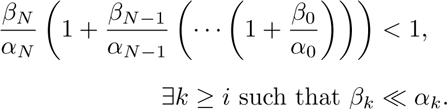

